## Supplemental Tables S1-S3 for "Refining the resolution of the yeast genotype-phenotype map using single-cell RNA-sequencing"

### SUPPLEMENTARY TABLES

| Regression predictor | *t* value | *p*-value |
|---|---|---|
| HMM-corrected genotype uncertainty | -9.89 | <2.00E-16 |
| Reference panel strain genome uncertainty | -8.36 | <2.00E-16 |
| Breadth of coverage | -2.66 | 8.04E-3 |
| Number of breakpoints | -2.00 | 4.60E-2 |
| Number of reads | -0.10 | 9.18E-1 |
| Depth of coverage | 0.08 | 9.41E-1 |

**Supplementary table 1 Predictors of the relatedness between high-coverage cells and their closest batch 1 lineage.** We used these variables as explanatory variables in a linear regression model where the relatedness of the high-coverage cells was the response variable. The high-coverage cells were defined as having a coverage in the top 25% of the distribution (>Q3). The data were normalized before performing the linear regression and the influence of the predictors is ranked in decreasing order from top to bottom of the table.

| Chromosome | QTL position | Effect size | KEGG gene annotation (60) |
|---|---|---|---|
| chr01 | 178191 | -0.00337928 | NA |
| chr02 | 469649 | -0.004014246 | YBR112C |
| chr03 | 271959 | 0.005822769 | YCR093W |
| chr04 | 129666 | -0.003667343 | YDL224C |
| chr04 | 277642 | 0.00314934 | YDL122W |
| chr04 | 551436 | -0.002390928 | NA |
| chr04 | 721119 | 0.003213192 | NA |
| chr05 | 195488 | 0.0041835 | YER020W |
| chr07 | 142006 | 0.004597008 | YGL197W |
| chr07 | 544909 | 0.003707906 | YGR032W |
| chr07 | 960936 | -0.0031564 | YGR234W |
| chr08 | 510386 | 0.000987114 | YHR188C |
| chr10 | 440709 | 0.004797933 | YJL005W |
| chr10 | 660787 | 0.013632744 | YJR127C |
| chr11 | 189721 | 0.003792844 | YKL109W |
| chr11 | 581166 | 0.004065964 | YLL061W |
| chr12 | 362424 | 0.005538545 | YLR115W |
| chr12 | 498456 | 0.011796208 | NA |
| chr12 | 647721 | 0.01117107 | NA |
| chr12 | 801095 | -0.002798822 | YLR309C |
| chr12 | 951031 | 0.00991076 | NA |
| chr13 | 51141 | 0.011146938 | YML120C |
| chr13 | 343663 | 0.009595804 | NA |
| chr14 | 442660 | 0.00010508 | YNL094W |
| chr14 | 478696 | 0.025678619 | YNL079C |
| chr15 | 88067 | 0.002939759 | YOL134C |
| chr15 | 163702 | -0.005493485 | YOL081W |
| chr15 | 471091 | -0.009577858 | NA |
| chr15 | 944240 | -0.004171259 | YOR316C |
| chr16 | 515135 | -0.002926068 | YPL023C |
| chr16 | 726411 | 0.002966014 | YPR084W |

**Supplementary table 2 QTL identified from the bulk fitness and DNA sequencing assays.**

| Chromosome | QTL position | Effect size | KEGG gene annotation (60) |
|---|---|---|---|
| chr01 | 37255 | 0.002989743 | YAL056W |
| chr02 | 509588 | -0.00570928 | YBR112C |
| chr03 | 204909 | 0.006294488 | NA |
| chr04 | 849797 | 0.002871261 | NA |
| chr04 | 1359610 | -0.003216782 | NA |
| chr05 | 189186 | 0.007163705 | YER020W |
| chr07 | 124719 | 0.004653679 | YGL197W |
| chr07 | 391249 | 0.004234537 | YGL071W |
| chr07 | 972982 | -0.002685337 | YGR234W |
| chr08 | 465706 | 0.00218292 | YHR188C |
| chr10 | 422419 | 0.007224158 | YJL005W |
| chr10 | 657711 | 0.01419846 | YJR127C |
| chr11 | 188878 | 0.005193796 | YKL109W |
| chr11 | 613622 | 0.004810808 | YLL061W |
| chr12 | 498456 | 0.016837383 | NA |
| chr12 | 591551 | -0.005087786 | YLR223C |
| chr12 | 657026 | 0.027398457 | NA |
| chr12 | 951087 | 0.012451375 | NA |
| chr13 | 50890 | 0.010534031 | YML120C |
| chr13 | 331755 | 0.008646562 | NA |
| chr14 | 314157 | 0.004021681 | YNL192W |
| chr14 | 481076 | 0.027917558 | YNL079C |
| chr15 | 73395 | 0.002153585 | YOL134C |
| chr15 | 194853 | -0.008589995 | YOL081W |
| chr15 | 467983 | -0.013085813 | NA |
| chr15 | 602954 | 0.002917765 | NA |
| chr15 | 992166 | -0.002155523 | YOR370C |
| chr16 | 511943 | -0.005435177 | YPL023C |
| chr16 | 720831 | 0.00578315 | YPR084W |

**Supplementary table 3 QTL identified from single cells HMM-corrected genotypes and closest lineage fitness.**
