## Supplemental Figures S1-S8 for "Refining the resolution of the yeast genotype-phenotype map using single-cell RNA-sequencing"

**SUPPLEMENTARY MATERIAL**

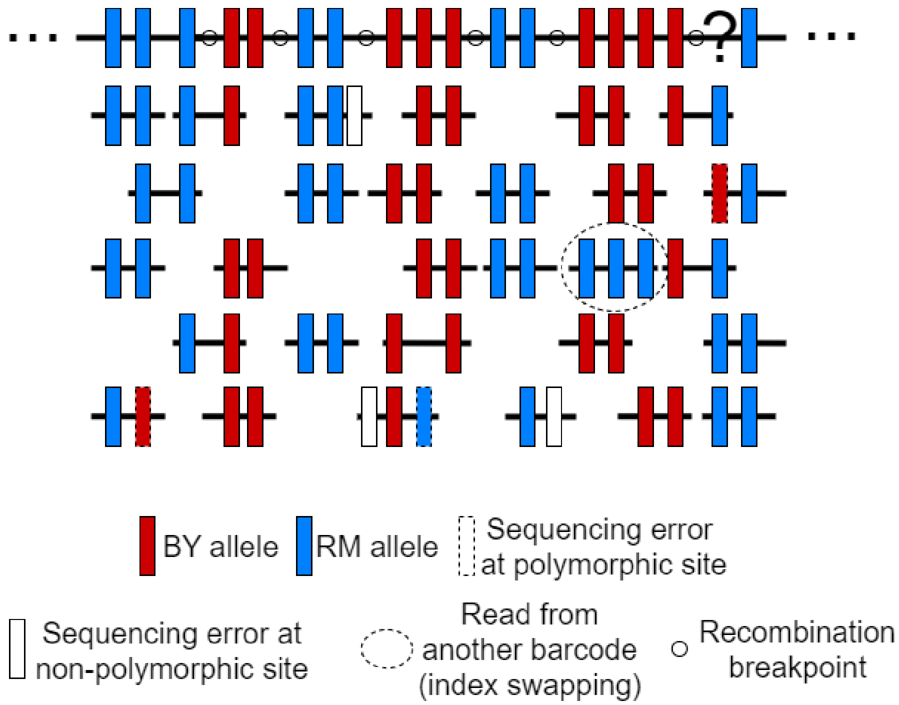


**Figure S1 Illustration of scRNA-seq reads used to estimate the HMM error rate parameters.** Reads from a single cell are being mapped to the RM parent reference sequence to obtain a reconstructed genome, i.e. the large black-filled rectangle at the top. The polymorphic sites represented on the reconstructed sequence either represent the RM allele (blue rectangle) or the BY allele (red rectangle). Although it is not possible to distinguish true sequencing errors at polymorphic sites (dashed line rectangle) from true mutations observed at a low frequency in a population, the sequencing error rate can be inferred by counting the proportion of non-polymorphic sites being mutated in the reads (white-filled rectangle). As for the rates at which reads come from other cells (index swapping), it can be inferred by the number of reads with at least 2 mismatches, i.e. a different allele than the rest of the reads mapping at common sites. After inferring the error rates, we can use the HMM to infer the most likely genotypes at each polymorphic site, e.g. the interrogation point is probably a RM allele as supported by the reads.

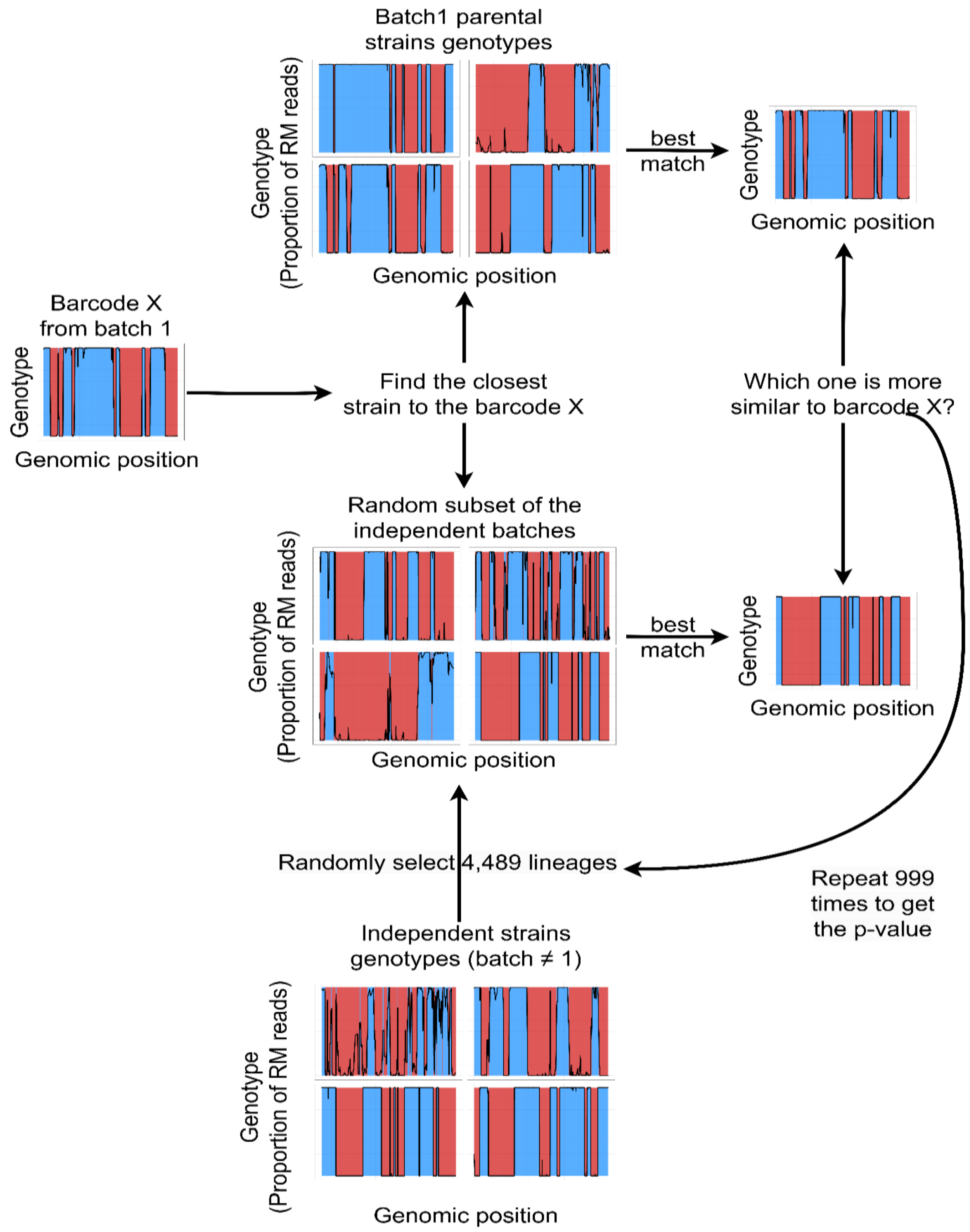


**Figure S2 Barcodes lineage assignment.** For every single cell barcode, the closest lineages in the batch 1 and in a random batch of the same size are selected based on the minimal expected distance. The random batch is generated by pooling 4,489 random lineages from the 22 batches that are independent from batch 1. Next, the distance between the barcode genotype and its best match in batch 1 is compared to the distance with the best match in the random batch. This procedure was repeated 999 times and the p-value was then estimated by the proportion of random subsamples from other batches that contained a better match to the single cell genotype than batch 1.

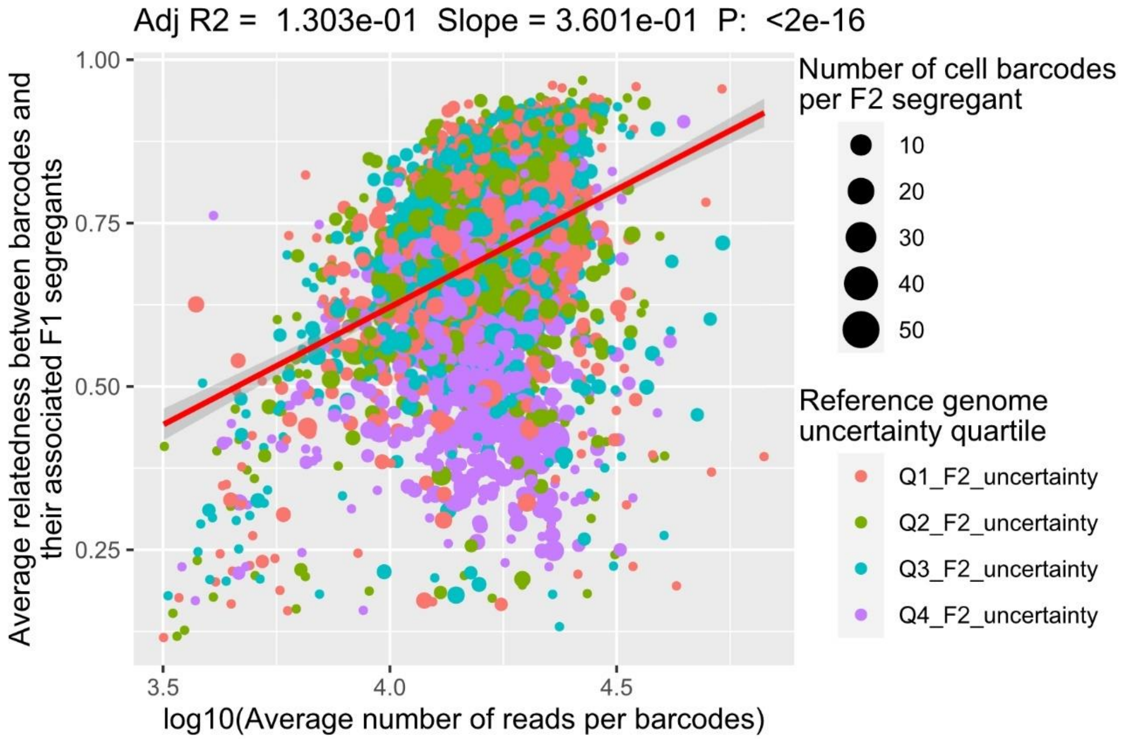


**Figure S3 Factors explaining the relatedness between scRNA-seq barcodes and their associated F2 segregant**. The relatedness between the barcodes and the F2 segregant is measured with the adjusted $R^2$ between the genotypes. The linear regression model is performed after a log10-transformation of the coverage (number of reads per barcode). Each dot represents a F2 segregant. The number of scRNA-seq barcodes associated to each segregant is represented by the dot size while its color represents the quartile of the F2 segregant genome uncertainty.

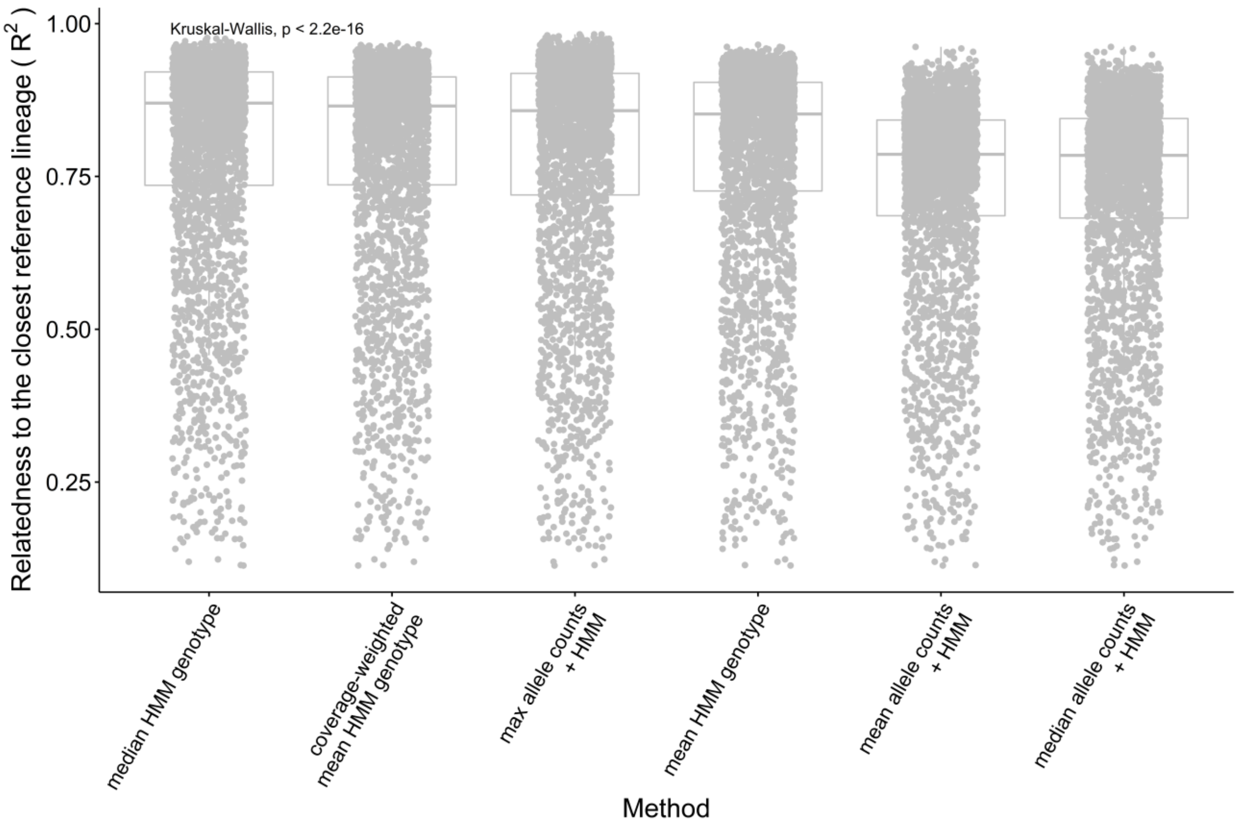


**Figure S4. Performance of the consensus genotype methods**. The relatedness between the consensus genotypes and the closest lineage in batch 1 is measured with the adjusted $R^2$ between the genotypes. The consensus genotype of each lineage is either obtained by calculating a summary statistic with the single-cell genotypes assigned to that lineage or by performing an HMM inference after combining the single cells allele counts using the summary statistics. The three summary statistics applied are the max, the mean and the median. The lineage assigned to a group of single cells is the one with the closest genotype across batch 1 F2 segregants.

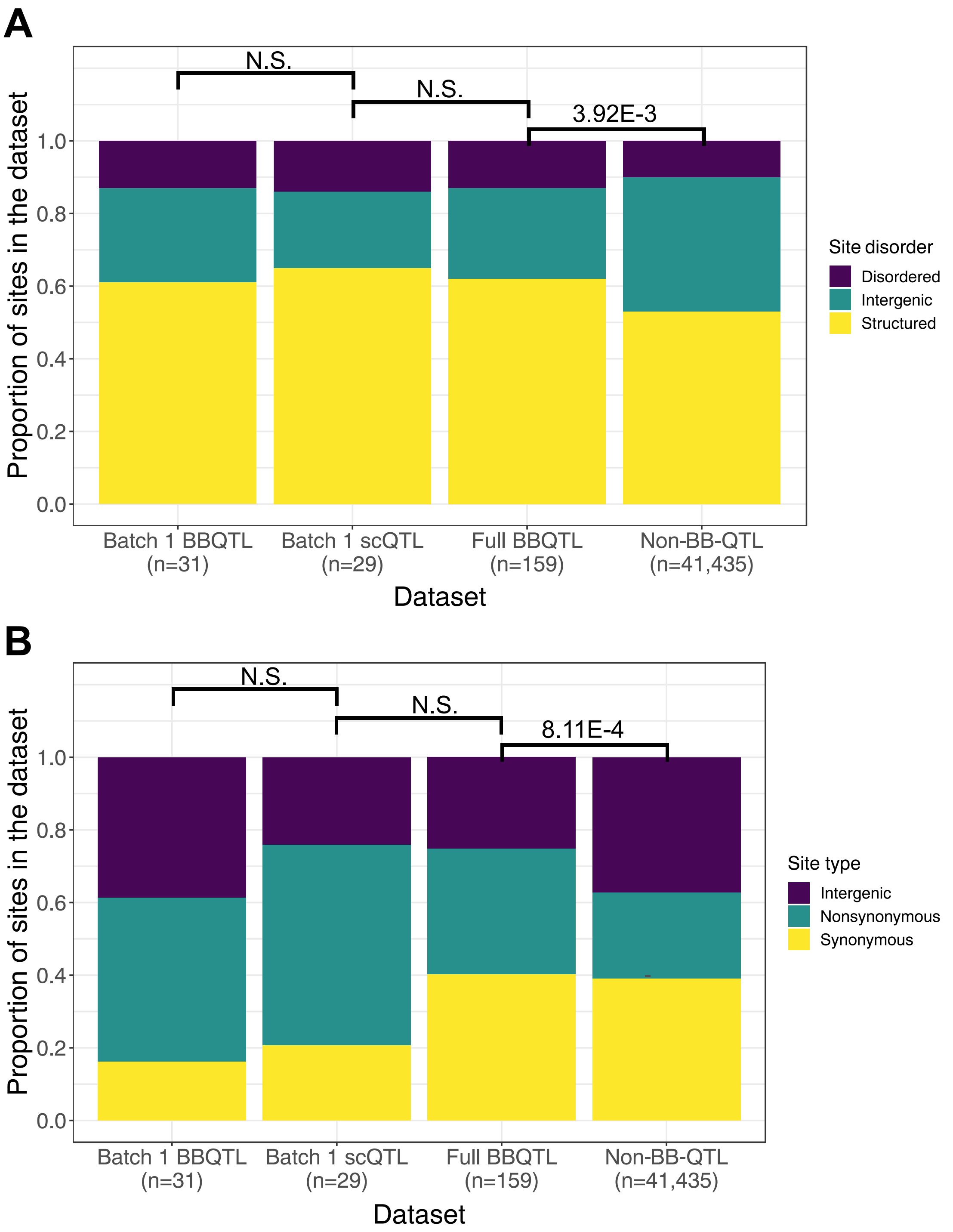


**Figure S5 Comparison of site features distribution across QTL models.** A) Site disorder distributions. The non-BB-QTL set represents polymorphic sites that are not identified as QTL in the model performed on the full dataset of ~100,000 segregants. The size of each set of polymorphic sites is indicated below its name on the x-axis. Fisher's exact test p-values are indicated above the boxplots where N.S. represents a p-value >=0.05. B) Site type distributions.

Then, I compared the distributions of these two site features in the full BB-QTL dataset, which are similar in the scRNA-seq, to the distributions in the full set of SNPs using Fisher's exact test and odd ratios. I also performed the phenotype variance partitioning based on the QTL features to identify which sites contribute the most to phenotype variations. Nonsynonymous sites have significantly higher odds of being QTL than intergenic sites. The trend is modest (odd ratio = 2.17; Fisher’s exact test p =1.94e-4) and the difference with synonymous sites is not significant. Furthermore, disordered sites have significantly higher odds of being QTL than intergenic sites although the trend is modest (odd ratio = 2.00; Fisher’s exact test p =1.67e-2) but the difference with structured sites is not significant.

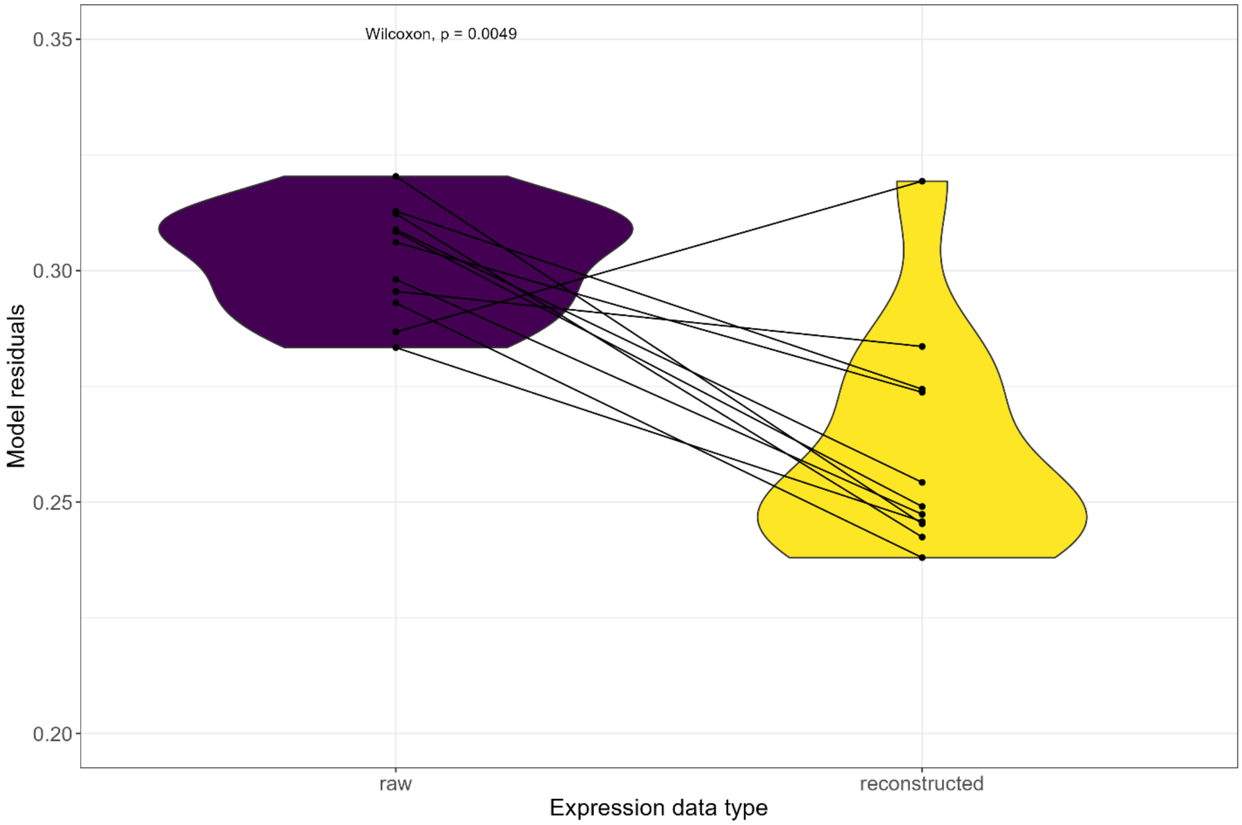


**Figure S6 Denoising the expression data with DISCERN decreases the residuals of the variance partitioning model.** We partitioned the phenotype variance based on the contribution of gene expression and genotype changes using the GREML method. The expression data were either the original ones obtained from our scRNA-seq assay (purple) or the reconstructed expression data obtained by denoising the raw expression data with DISCERN (yellow). Each point pair represents a sample size, and the Wilcoxon signed rank p-value is indicated at the top of the graph.

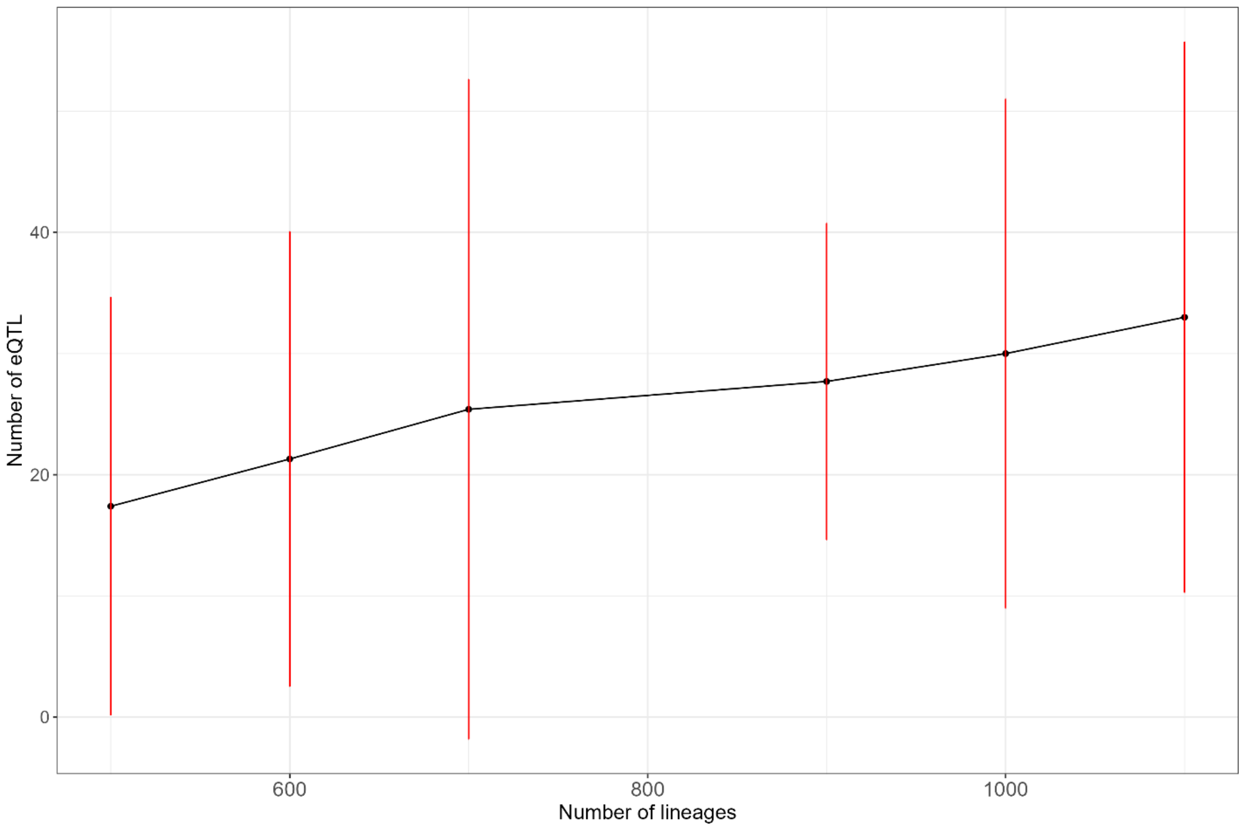


**Figure S7 The number of detected eQTL revealed by scRNA-seq increases with sample size.** We evaluated the number of eQTL detected per gene for the 10 most heritable genes in our dataset. For each sample size, the dot represents the average number of eQTL detected and the red line represents 2 standard deviations.

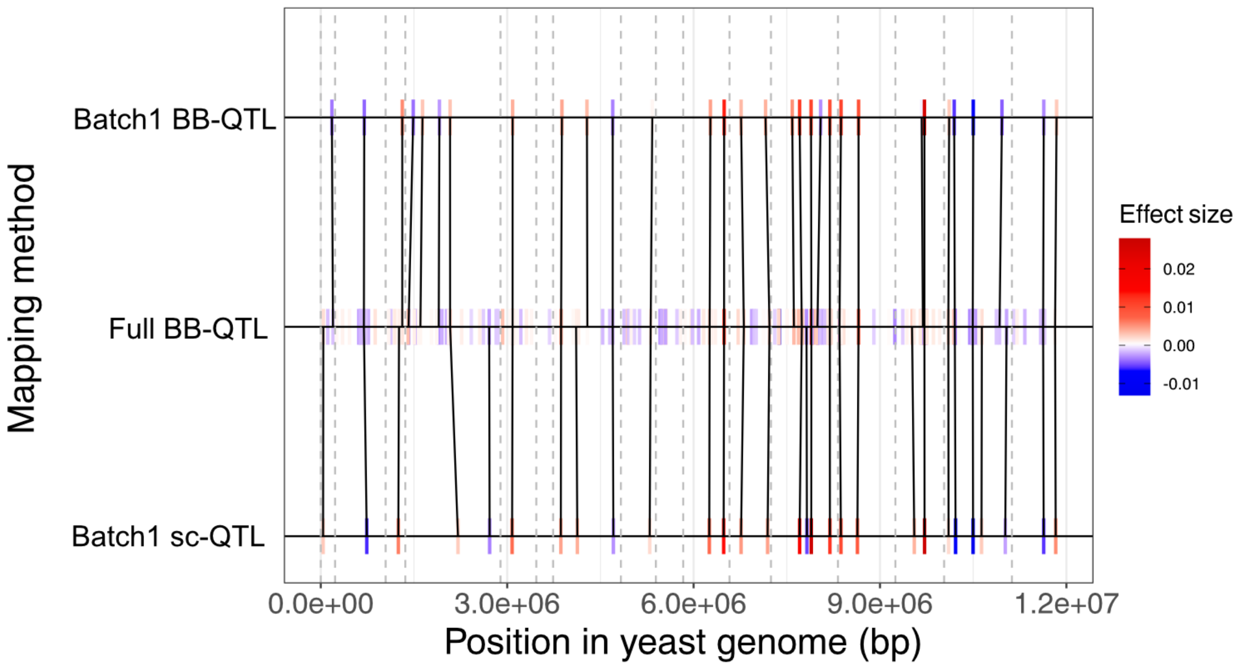


**Figure S8 Comparison of the QTL mapping models for the 30C phenotype**. The black lines connect the QTL that are shared between 2 models. These shared QTL are identified by the Needleman-Wunsch algorithm which is based on the minimization of recombination distance, effect size difference and allele frequency weighting as described by Nguyen Ba et al., 2022. QTL effect size is represented by a color gradient where negative effects are represented in blue while positive effects are represented in red. The grey dashed lines represent the start position of each of the 16 chromosomes in the yeast genome.
